# Astrocytic VMAT2 in the developing prefrontal cortex is required for normal grooming behavior in mice

**DOI:** 10.1101/2021.01.27.428434

**Authors:** Francesco Petrelli, Tamara Zehnder, Luca Pucci, Corrado Cali, Bianca Maria Bondiolotti, Alicia Molinero Perez, Glenn Dallerac, Nicole Déglon, Bruno Giros, Fulvio Magara, Lorenzo Magrassi, Jean-Pierre Mothet, Linda Simmler, Paola Bezzi

## Abstract

Astrocytes control synaptic activity by modulating peri-synaptic concentrations of ion and neurotransmitters including dopamine and, as such, can be critically involved in the modulation of some aspect of mammalian behavior. Here we report that genetic mouse model with a reduced medial prefrontal cortex (mPFC) dopamine levels, arising from astrocyte-specific conditional deletion of vesicular monoamine transporter 2 (VMAT2; aVMTA2cKO mice) shows excessive grooming and anxiety-like behaviour. The VMAT2cKO mice also develop a synaptic pathology, expressed through increased relative AMPA vs. NMDA receptor currents in synapses of the dorsal striatum receiving inputs from the mPFC. Importantly, behavioural and synaptic phenotypes are prevented by reexpression of mPFC VMAT2, showing that the deficits are driven by mPFC astrocytes. By analysing human tissue samples, we found that VMAT2 is expressed in human mPFC astrocytes, corroborating the potential translational relevance of our observations in mice. Our study shows that impairments of the astrocytic-control of dopamine in the mPFC has a profound impact on circuit function and behaviours, which resemble symptoms of anxiety disorders and obsessive compulsive disorder (OCD).

## Introduction

Self-grooming is an innate behaviour that most animal species (including humans) practise in order to maintain bodily hygiene (1), but it can become pathological under conditions of stress or in the case of neuropsychiatric disorders (2–4), such as obsessive-compulsive disorder (OCD) and other OC-spectrum disorders, characterised by excessive repetitive behaviours (5). OCD is a highly heritable (h 2 = 0.27–0.65) (6), heterogeneous and debilitating neuropsychiatric disorder that affects 2–3% of the population worldwide (7–10) and is associated with persistent intrusive thoughts (obsession), repetitive rituals (compulsion), and excessive anxiety that significantly impairs social activities. The heterogeneity of OCD is related to the various types of obsessions, compulsions and co-morbid conditions associated with it (11). In addition, there are other neuropsychiatric OC-spectrum disorders such as Tourette’s syndrome and trichotillomania that share features of OCD, particularly the occurrence of repetitive maladaptive behaviours.

The neurobiological basis of repetitive behaviours in OCD is still unclear, especially in terms of the neural circuits and neuromodulators involved. Convergent neuroimaging and neurocognitive findings support a model in which abnormal cortico-striato-thalamo-cortical (CSTC) loops play a critical role in the symptoms (12,13). These findings have been confirmed by studies of genetically engineered mouse lines (14–16), which have characterised the CSTC loop dysfunctions implicated in OCD pathophysiology in detail and provided invaluable insights into the relationships between striatal (ST) dysfunctions and abnormal OCD-relevant behaviours (14,17). It is still unknown whether and how upstream cortical structures are involved in generating the striatal abnormalities leading to pathological behaviours. However, it seems that prefrontal cortex (PFC) dysfunctions are central in the CSTC model as patients show orbitofrontal cortex (OFC) hyperactivity at baseline and after symptom provocation (18–20), and the PFC (and striatal) hyperactivity correlates with the severity of the obsessions, compulsions and symptom-associated affective states (21). Moreover, the persistence of maladaptive patterns of inflexible thoughts and behaviours in patients (22,23) and in mouse models of OCD (24–26) indicate a lack of cognitive flexibility (i.e. the ability of the PFC to adapt behaviours in response to changing situational requirements). Preclinical studies of mouse PFC have confirmed its possible role in the pathophysiology of OCD (25,27–31), and one study of how orbitofrontal-striatal projections contribute to OCD-relevant repetitive behaviour has shown that chronic optogenetic stimulation in normal mice is sufficient to induce excessive grooming (31). However, despite these advances, there are fewer published studies of the PFC in the context of OCD than those of striatal abnormalities.

Over the last 20 years, studies of the neuromodulators involved in OCD have mainly concentrated on serotonergic dysfunctions because it has been shown that serotonin reuptake inhibitors (SRIs) are effective in treating the repetitive and compulsive behaviour (32–38). However, nearly half of OCD patients do not sufficiently respond to SRIs treatment (39), and it seems that the dopaminergic (DAergic) pathways are also involved in the pathogenesis of OCD (40–44), as antipsychotics as add-on medication have proved to be clinically beneficial in OCD patients who do not respond to SRIs (45–47), and microdialysis studies of rodents have shown that antipsychotics increase DA levels in the PFC (48,49). The involvement of DAergic neuromodulation in the pathophysiology of the repetitive compulsive behaviours characterising OCD spectrum disorders is particularly intriguing as functional neuroimaging studies have revealed that OCD patients show abnormal hyperactivity in the PFC, where DAergic innervation is particularly abundant. However, despite the importance of the PFC and DA in inducing or aggravating the repetitive and compulsive behaviour characterising OCD, we still largely ignore the possible role of PFC DAergic dysfunctions in the pathophysiology of OCD.

In this context, it is particularly interesting to note that PFC astrocytes are endowed with the unique features of DAergic glial cells, and are responsible for regulating DA homeostasis (50). The astrocytic regulation of DA occurs in the developing PFC, and depends on the expression of vesicular monoamine transporter 2 (VMAT2), an intracellular transporter known to be involved in regulating DA homeostasis in various monoaminergic cell types (51). Astrocyte VMAT2 controls monoamine oxidase B (MAOB)-dependent metabolic capacity by sequestering DA from the cytoplasm, and mice in which astrocyte VMAT2 has been conditionally deleted show increased MAOB activity and a consequently significant decrease in extracellular DA levels. The unbalanced DA levels in the PFC of mice lacking astrocyte VMAT2 has been associated with increased basal activity in PFC circuits and deficits in behavioural flexibility (50), a phenotype that is reminiscent of some of the physiological and cognitive aspects of OCD patients and mouse models of OCD (22).

The findings of this study show that VMAT2 is extensively expressed in astrocytes located in the frontal cortical tissue of mice and humans, and that mice carrying the medial PFC (mPFC) astrocyte VMAT2 deletion have a behavioural phenotype that is similar to that of mouse models of OCD: excessive grooming and increased anxiety. These behaviours are rescued by the selective re-expression of VMAT2 in PFC astrocytes and by long-term L-DOPA treatment, thus indicating that astrocyte VMAT2 and appropriate DA levels prevent the onset of the OCD-like behaviour of aVMAT2cKO mice. Furthermore, the role of astrocyte VMAT2 in neuronal pathology is further highlighted by optogenetic projection targeting, showing that the absence of VMAT2 in mPFC astrocytes can modulate mPFC corticostriatal synaptic strength, part of the circuitry implicated in OCD.

## Results

### Loss of VMAT2 in astrocytes leads to a behavioural phenotype related to obsessive compulsive disorder

In order to study the physiological significance of astrocytic VMAT2 in OCD-relevant phenotypes, we used our previously reported inducible knock-out mouse line (50) in which the protein can be specifically deleted in a temporally controlled manner. The mouse line is a crossing of mice harbouring the tamoxifen(TAM)-inducible *cre* recombinase transgene driven by the human astrocytic glial fibrillary acidic protein (hGFAPcre) promoter (52) with mice containing *cre*-excisable *loxP* sequences in the endogenous *VMAT2* gene (VMAT2^loxP/loxP^) (53). The progeny that inherited both alleles and were treated with TAM from P20 to P28 presented the selective deletion of astrocyte VMAT2, and are here referred to as aVMAT2cKO mice; the controls were VMAT2^loxP/loxP^ mice injected with TAM (control LoxTAM). The specificity and efficacy of the TAM-induced astrocyte VMAT2 excision was confirmed in the fluorescent Cre^ERT2^XR26-tdTomato-reporter (54) and aVMAT2cKO mice lines. Confocal microscopy revealed the expression of tdTomato in the astrocytes of different brain areas, and showed recombination in 60% of PFC astrocytes as previously reported (54); no recombination was detected in cortical neurons.

Anatomical and histological brain analyses showed that they were grossly normal (50) but, starting at 4-5 months of age, they developed facial hair loss (Figure 1a). The penetrance of this phenotype increased with age and affected most (90%) of the transgenic mice, which developed hair loss regardless of whether they were housed alone or with cage mates. The videotaping of habituated, individually housed aVMAT2cKO mice, showed that they engaged in more grooming bouts and spent significantly more time self-grooming than their littermates (Figure 1b, 1c, 1d), whereas their digging and jumping activities were normal, thus indicating a possible susceptibility to the development of self-directed stereotypies (Figure 1b) (55).

**Figure 1.**
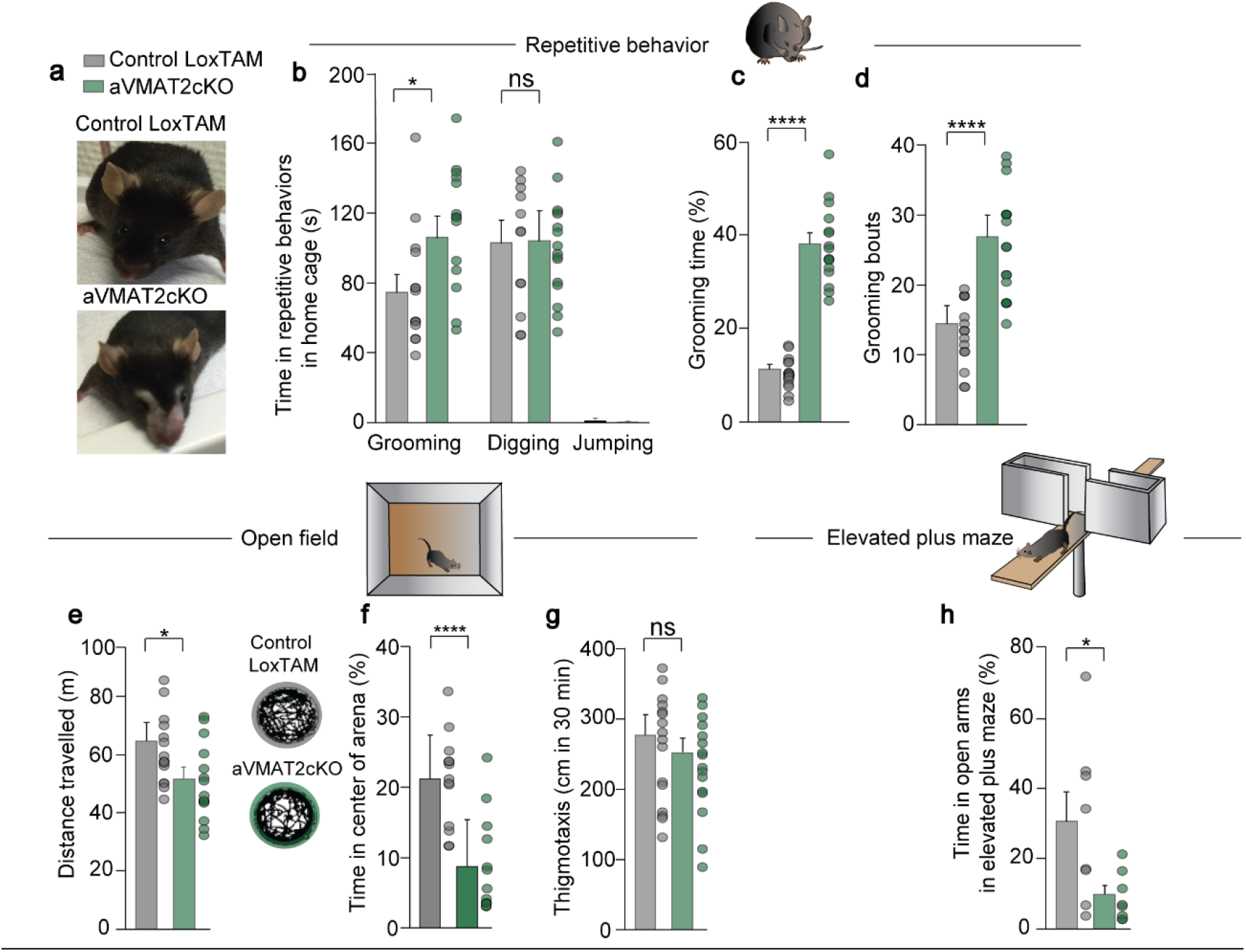
Facial lesions, excessive grooming and anxiety-like behaviours in aVMAT2cKO mice. **a**) aVMAT2cKO mice that have removed hair from facial regions without creating large lesions. **b-d**) Repetitive behavioural tasks. **b**) The histograms show the average time that control LoxTAM (grey) and aVMAT2cKO mice (green) spent in grooming, digging and jumping. The error bars indicate the SEM *p<0.005 (n=16 mice in each group; unpaired Student’s t-test). **c**) The graph shows the percentage of time that the control LoxTAM and aVMAT2cKO mice spent in grooming ****p<0.0001 (n=15 in each group; unpaired Student’s t-test). The error bars indicate the SEM. **d**) The graph shows the grooming bouts of the control LoxTAM and aVMAT2cKO mice ****p<0.0001 (n=15 in each group; unpaired Student’s t-test). The error bars indicate the SEM. **e-g**) Open field tasks. The histograms show the average total distance travelled by control LoxTAM and aVMAT2cKO mice during 10 minutes of free exploration of a maze (**e**), and the average percentage of time that control LoxTAM and aVMAT2cKO mice spent (**f**) in the centre of the arena, and (**g**) thigmotaxis (cm) over 30 minutes *p<0.01; ****p<0.0001 (n=10-16 in each group; unpaired Student’s t-test). The error bars indicate the SEM. **h**) Elevated plus maze tasks. The histograms show the average percentage of time that the control LoxTAM and aVMAT2cKO mice spent in the open arms. The error bars indicate the SEM.*p<0.01 (n=8 in each group unpaired Student’s t-test).

As the aVMAT2cKO mice showed compulsive-like behaviour, we used open-field and elevated plus maze tests to examine anxiety-like behaviour in order to investigate whether their phenotypes further resembled OCD spectrum disorders. A novel open-field task showed that the mice engaged in less locomotor activity (Figure 1e) and spent much less time exploring the central area than their control littermates (Figure 1f), which is typical of mice with anxiety-like phenotypes. Furthermore, there was no difference between control and aVMAT2cKO mice in terms of activities along the walls and in the corners, areas that are believed to be less stressful (Figure 1g). These signs of anxiety were confirmed by the elevated plus maze test, which showed that aVMAT2cKO mice took longer to cross into the open areas (a riskier environment) and spent less time exploring the open than the closed arms of the maze (Figure 1h). Taken together, these observations indicate that aVMAT2cKO mice are characterised by excessive and compulsive self-grooming behaviour associated with increased anxiety, which can be considered analogous to the pathological repetitive behaviours associated with OCD spectrum disorders (56,57).

### Loss of VMAT2 in astrocytes is associated with a strengthening of PFC-striatum transmission in MSNs

The OCD-relevant repetitive behaviours and anxiety, and the significant increase in the basal activity of the mPFC observed in aVMAT2cKO mice (50), together with findings of long-lasting changes in cortical-striatal neurotransmission in animal models of compulsive behaviour (including animal models of OCD) (58), suggested that the mPFC-dorsomedial striatum (DMS) pathway (59) may be involved in the compulsiveness of aVMAT2cKO mice.

We injected an adeno-associated virus (AAV8-Syn-ChrimsonR-GFP) in the mPFC that anterogradely labelled fibre terminals in the medial part of the dorsal striatum as previously reported (59) (Figure 2a). Then, we measured the ratio between AMPA and NMDA receptor excitatory postsynaptic currents (EPSCs) as a proxy of excitatory synaptic strength in dorsomedial putative medium-sized spiny neurons (MSNs), using channelrhodopsin (ChR2) ChrimsonR (60) expression in the mPFC to activate cortical fibres specifically (Figure 2c). The PFC was bilaterally injected with adeno-associated virus (AAV8) to express a fusion protein of a red-shifted ChR2 and enhanced green fluorescent protein (ChrimsonR-GFP) under the Syn promoter (60,61) in order to target cortical pyramidal neurons of control and VMAT2cKO mutant mice (Figure 2a). Pulses of orange LED light (593 nm, 1 ms pulses at 0.1 Hz) were delivered through the patch-clamp microscope objective to the mPFC terminals in striatum slices and the EPSCs in the putative MSNs in the DMS were simultaneously recorded (Figure 2b).

**Figure 2.**
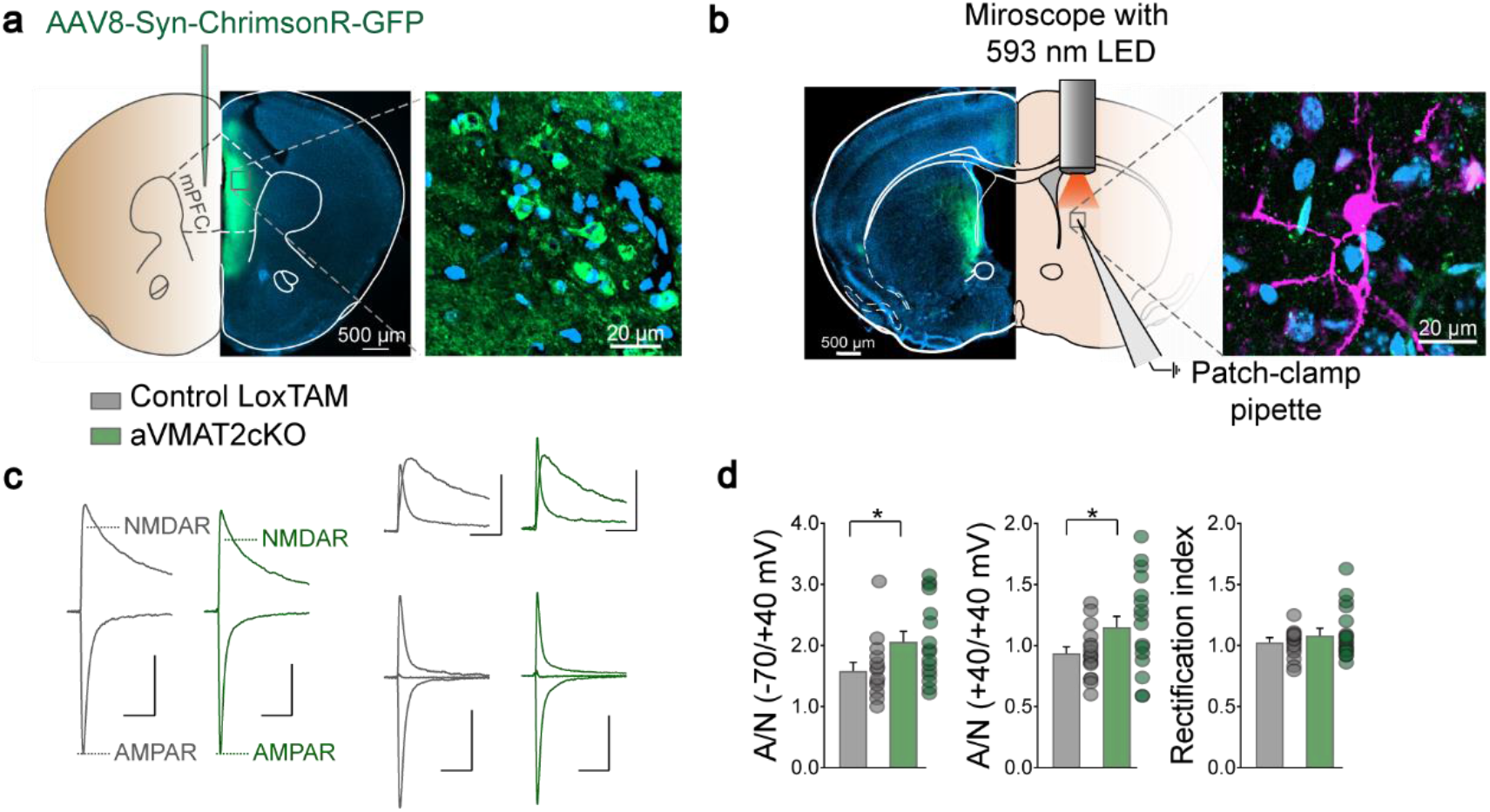
Alterations in mPFC-to-DMS synapses of aVMAT2cKO mice. **a**) Location of virus injection and infection. AAV8-Syn-ChrimsonR-GFP was injected into the mPFC of the control LoxTAM and aVMAT2cKO mice. Cell nuclei were stained with Hoechst (blue). **b**) Whole-cell patch clamp recordings of striatal slices. The terminals of the mPFC-to-dorsomedial striatum (DMS) projections were stimulated with orange light (593 nm), and the light-evoked currents were recorded in putative medium-size spiny neurons (MSNs). For illustrative purposes, a patch-clamped MSN was filled with biocytin (pink in right-hand picture). **c**) Example traces of patch-clamp recordings from LoxTAM (grey) and aVMAT2cKO mice (green): left: A/N (−70/+40 mV); top right: A/N (+40/+40 mV); bottom right: rectification index. **d**) A/N ratios at holding currents of −70/+40 mV (left panel; n (cells) = 18 LoxTAM, 19 aVMAT2cKO; Mann-Whitney test, *p < 0.05) and +40/+40 mV (middle panel; n (cells) = 17 LoxTAM, 20 aVMAT2cKO; unpaired t-test: *p <0.05), were significantly increased in the aVMAT2cKO mice, but the rectification index (right panel) was not significantly different between genotypes (n (cells) = 18 LoxTAM, 20 aVMAT2cKO; Mann-Whitney test: p = 0.89). Data are mean ± SEM.

Evaluation of the changes in the AMPAR: NMDAR current ratio (A/N ratio) in putative MSNs in the DMS showed that it was increased in the aVMAT2cKO mice (Figure 2c and 2d), indicative of an increase in synaptic strength. An assessment of AMPAR-mediated currents at negative (−70 mV), reversal (0 mV), and positive potentials (+40 mV) in order to test for the presence of AMPARs lacking GluA2 in the same preparation gave a rectification index (RI) of 1.03 in the MSNs of the control mice and 1.09 in those of the aVMAT2cKO mice (Figure 2d), thus slightly more AMPARs lacking GluA2 were found in the absence of astrocyte VMAT2. Taken together, these findings indicate that the loss of VMAT2 in astrocytes is associated with strengthening of PFC transmission to DMS MSNs.

### Restoration of VMAT2 in mPFC astrocytes rescues OCD-relevant phenotypes

To determine the significance of VMAT2 and the associated decrease in DA in the developing mPFC for the expression of OCD-relevant phenotypes, we evaluated the effects of VMAT2 re-expression in mPFC astrocytes on the increased self-grooming and anxiety-like behaviours of aVMAT2cKO mice. As shown in Figure 1, aVMATcKO mice spent significantly more time grooming in the open field than control mice. A single intracranial injection of the astrocyte-specific lentiviral vector (LentiVMAT2) (50,62) at P25 (Figure 3a and 3b), which enables the selective re-expression of VMAT2 in astrocytes and normalized basal extracellular DA levels in the mPFC (50), reduced the level of aVMAT2cKO grooming activity (Figure 3c, 3d, 3e). In contrast, the control lentiviral injection (LentiGFP) did not modify the basal grooming behaviour of the control mice. In line with our previous observations, the aVMAT2cKO mice spent significantly less time in the centre of the open field than their control littermates (Figure 3h), thus indicating an anxiety-like phenotype. The injection of LentiVMAT2 corrected this phenotype but control LentiGFP did not significantly affect the centre time of the control mice (Figure 3h). LentiVMAT2 treatment also increased the open field locomotion activities of the aVMAT2cKO mice but not the control mice (Figure 3g). We further investigated the anxiolytic effects of VMAT2 using the elevated plus maze test. LentiVMAT2-injected aVMAT2KO mice spent more time in the open arms of the maze than the control-injected aVMAT2cKO mice (Figure 3f). Control lentiviral injection had no significant effects on any measure of anxiety in control mice, thus demonstrating that VMAT2 activity contributes to the heightened anxiety-like phenotypes of aVMAT2cKO mice.

**Figure 3.**
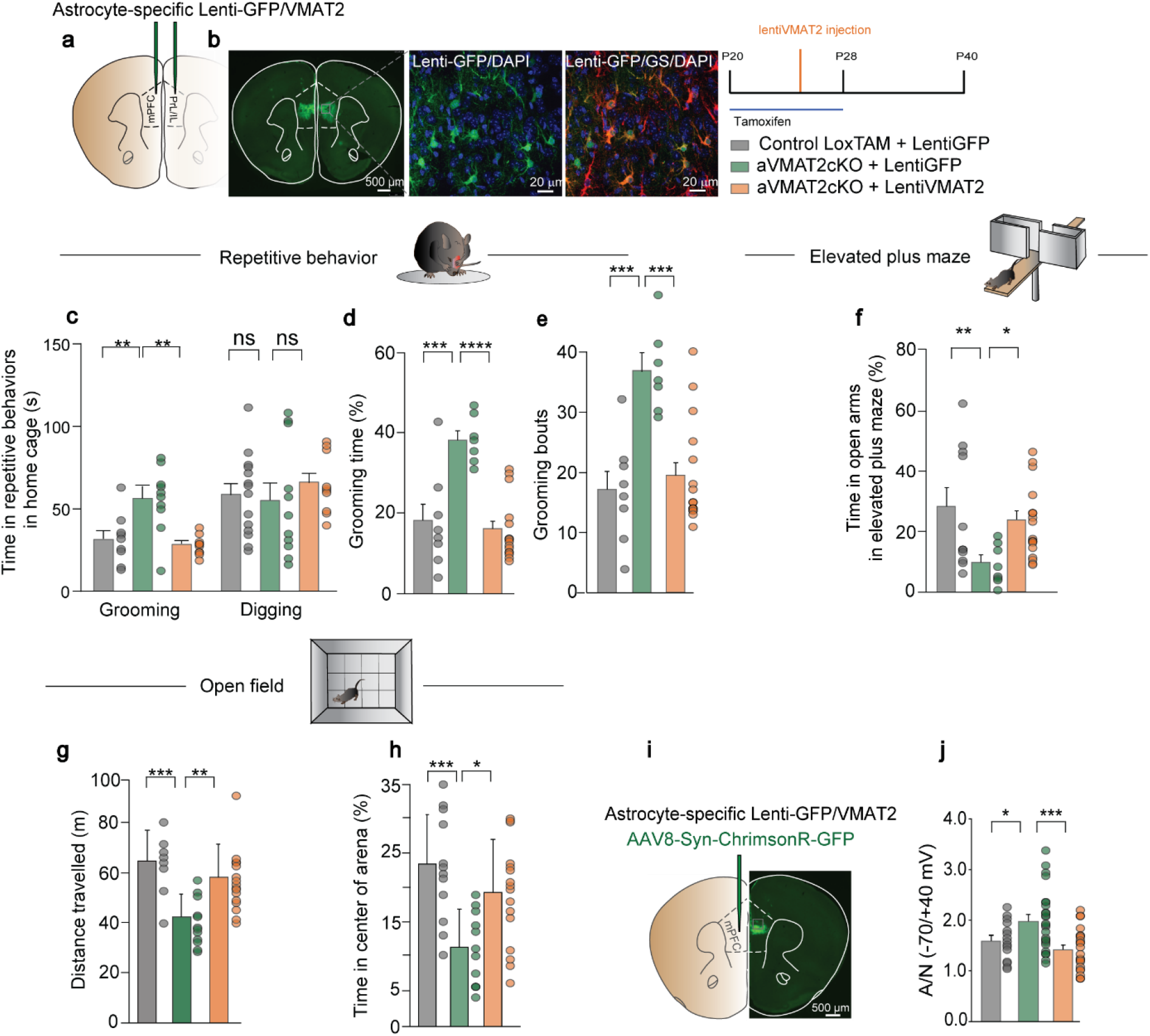
Astrocytic VMAT2 re-expression in the mPFC rescues the behavioural deficits and synaptic alterations of aVMAT2cKO mice. **a**) Schematic representation of the control GFP virus (LentiGFP) or LentiVMAT2 virus injection in the mPFC (prelimbic/infralimbic). **b**) Left: representative confocal coronal image showing the infection in the mPFC. Scale bar = 500μm. Centre: high magnification confocal images of LentiGFP (green), DAPI (blue) and glutamine synthase (GS) (red) in the mPFC. Scale bar = 20μm. Right: event timeline of intra-peritoneal injections of tamoxifen (P20-P28) and local infection with LentiVMAT2 or LentiGFP (P25). Behavioural and patch-clamp experiment were conducted starting P40. **c-e**) Repetitive behavioural tasks. (**c**) Average time that control LoxTAM mic (grey), aVMAT2cKO mice infected with LentiGFP virus (green) and aVMAT2cKO mice infected with LentiVMAT2 virus (orange) spent in grooming and digging. **p<0.005 (n=8 each group; one-way ANOVA, followed by Tukey’s *post hoc* HSD test) (**d**) Percentage of time that the control LoxTAM and aVMAT2cKO mice infected with LentiGFP (grey and green) or lentiVMAT2 virus (orange) spent in grooming ***p<0.001, ****p<0.0001 (n=8-10; one-way ANOVA, followed by Tukey’s *post hoc* HSD test). (**e**) The graph shows the grooming bouts of control LoxTAM and aVMAT2cKO mice infected with LentiGFP (grey and green) or lentiVMAT2 virus (orange) ***p<0.001 (n=8 each group; one-way ANOVA, followed by Tukey’s*post hoc* HSD test). **f**) Elevated plus maze tasks. The histograms show the average percentage of time that control LoxTAM and aVMAT2cKO mice infected with LentiGFP virus (grey and green) or lentiVMAT2 virus (orange) spent in the open arms. *p<0.01, **p<0.005 (n=10-14; one-way ANOVA, followed by Tukey’s *post hoc* HSD test) **g-h**) Open field tasks. (**g**) The histograms show the average total distance travelled by control LoxTAM and aVMAT2cKO mice infected with LentiGFP (grey and green) or lentiVMAT2 virus (orange) during 10 minutes of free exploration of a maze, and (**h**) the average percentage of time they spent in the center of the arena. *p<0.05, **p<0.005, ***p<0.001 (n=9-12 in each group, one-way ANOVA, followed by Tukey’s *post hoc* HSD test). **i-j**) Ex vivo slice electrophysiology. (**i**) Schematic of virus injection. (**j**) The aVMAT2cKO mice injected with lentiGFP (green) showed a significantly higher A/N ratio (measured at holding currents −70/+40 mV) compared to the LoxTAM + lentiGFP control group (grey), but the selective re-expression of VMAT2 in the mPFC of aVMAT2cKO mice (orange) prevented the emergence of the synaptic phenotype (n (cells)= 15 LoxTAM + lentiGFP, 25 aVMAT2cKO + lentiGFP, 22 aVMAT2cKO + lentiVMAT2; one-way ANOVA: P = 0.0009, F(2,59) = 7.948; Tukey’s*post hoc* test: *p <0.05, ***p <0.001)(J). The error bars indicate SEM.

In order to evaluate whether restoring DA levels in the mPFC during the period of mPFC maturation was effective in reducing the abnormal behaviour, the behavioural profiles of the aVMAT2cKO mice were subsequently evaluated after chronic systemic treatment with L-DOPA (20mg/kg^-^1 i.p.) or vehicle from P20 to P40. In line with the results obtained using LentiVMAT2, the increased anxiety and self-grooming significantly improved (Supplementary Figure 3).

We further examined whether the re-insertion of VMAT2 in astrocytes during postnatal mPFC development contributes to preventing the plasticity of the MSNs synapses of aVMAT2cKO mice (Figure 2) by assessing the A/N ratio at mPFC to DMS synapses. Control lentiviral injection had no significant effect, but LentiVMAT2 injection significantly prevented the increase in the A/N ratio in aVMAT2cKO mice (Figure 3i and 3j).

### Deletion of VMAT2 in mPFC astrocytes is sufficient to induce OCD-relevant phenotypes

In order to assess the sufficiency of mPFC aVMAT2 deletion to induce OCD-like behavioural phenotypes, we specifically deleted astrocytic VMAT2 in the mPFC by injecting VMAT2^loxP/loxP^ mice with a Lentiviral vector (62) carrying cre recombinase or GFP (53) on P20 (Figures 4a and 4b).

**Figure 4.**
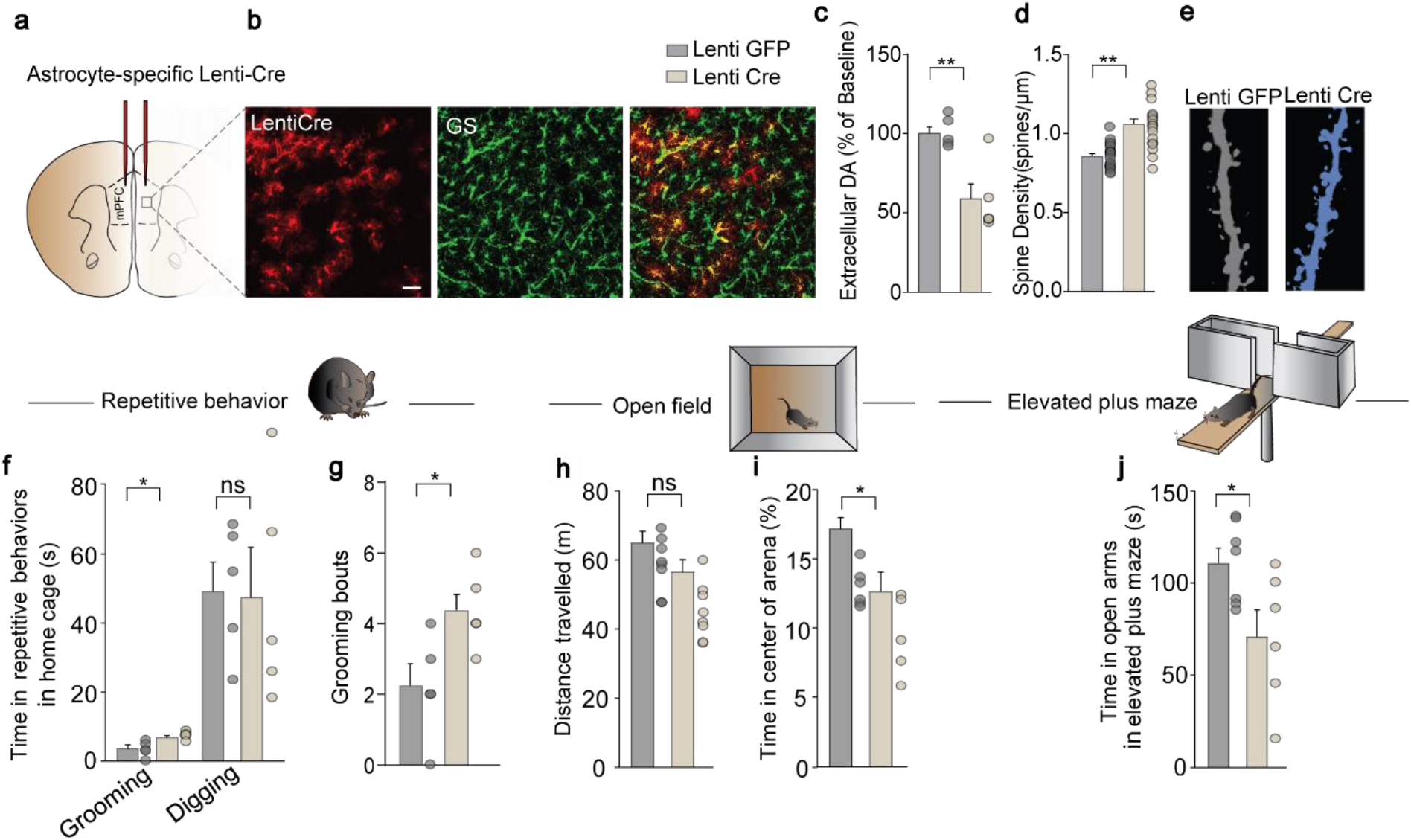
The deletion of VMAT2 in mPFC astrocytes is sufficient to cause OCD-relevant behavioural deficits. **a**) Schematic representation of Lentiviral vector carrying cre recombinase (LentiCre) or GFP (LentiGFP) injection in the mPFC (prelimbic/infralimbic). **b**) Representative high-magnification confocal images of LentiCre (red) and GS (green) in the mPFC. Scale bar = 50 μm **c**) Histograms show average DA levels calculated in extra-cellular perfusates of *in vivo* microdialysis in the PFC of control VMAT2^loxp/loxp^ mice infected with LentiGFP (grey) and VMAT2^loxp/loxp^ mice infected with LentiCre virus (beige), expressed as fold percentages of baseline levels. **p<0.01 (n=5 each group, unpaired Student’s t-test). The error bars indicate the SEM. **d**) Histograms of spine density calculated in layer V of the PFC of control LoxTAM-ThyEGFP mice and VMAT2^loxp/loxp^-ThyEGFP mice infected with LentiCre virus. **p<0.01 (n=16 each group, unpaired Student’s t-test). The error bars indicate the SEM. **e**) Representative confocal images showing Layer V mPFC dendritic spines in control LoxTAM-ThyEGFP mice and VMAT2^loxp/loxp^ –ThyEGFP mice infected with LentiCre virus. **f-g**) Repetitive behavioural tasks: (**f**) the histograms show the average time that control VMAT2^loxp/loxp^ mice infected with LentiGFP (grey) and VMAT2^loxp/loxp^ mice infected with LentiCre virus (beige) spent in grooming and digging. *p<0.05 (n=5 each group, unpaired Student’s t-test).The error bars indicate the SEM. (**g**) The graph shows the grooming bouts of VMAT2^loxp/loxp^ mice infected with LentiGFP (grey) and VMAT2^loxp/loxp^ mice infected with LentiCre virus (beige). *p<0.05 (n=5 each group, unpaired Student’s t-test). The error bars indicate the SEM. **h-i**) Open field tasks. (**h**) The histograms show the average total distance travelled by VMAT2^loxp/loxp^ mice infected with LentiGFP and VMAT2^lox/ploxp^ mice infected with LentiCre virus (beige) during 10 minutes of free exploration of a maze (n=5 each group, unpaired Student’s t-test), and (**i**) the average percentage of time they spent in the centre of the arena. *p<0.05 (n=5 each group, unpaired Student’s t-test).The error bars indicate the SEM. **j**) Elevated plus maze. The histograms show the average percentage of time that VMAT2^loxp/loxp^ mice infected with LentiGFP (grey) and VMAT2^loxp/loxp^ mice infected with LentiCre virus (beige) spent in the open arms. *p<0.05 (n=5 each group, unpaired Student’s t-test).The error bars indicate the SEM.

A single injection of LentiCre on P20 (Figure 4) induced phenotypes similar to those previously reported in aVMAT2cKO mice in whom VMAT2 was deleted from all astrocytes (50): decreased VMAT2 and extracellular levels of DA in the mPFC (Figure 4c), and increased spine density on P40 (Figure 4d and e). At behavioural level, LentiCre in the mPFC robustly increased the grooming activities of VMAT2^loxP/loxP^ mice, which were significantly greater than those of the VMAT2^loxP/loxP^ mice injected with LentiGFP virus (Figure 4f and 4g). A similar trend was observed in the anxious behaviour measured in the open field test and elevated plus maze (Figure 4h and 4i). In particular, LentiCre injection reproduced the anxiety-like phenotype of reduced open field centre time (Figure 4i) but not the decrease in distance travelled (Figure 4h). In the elevated plus maze, astrocyte-specific LentiCre injected in the mPFC had significant effects on other anxiety measures: it decreased the percentage of time spent in the open arms (Figure 4j). In brief, the results of these experiments showed that the specific deletion of VMAT2 from astrocytes in normal mice was sufficient to reproduce the core symptoms of grooming and anxiety-like behaviours that characterise the persistent OCD-like phenotype of the aVMAT2cKO mice.

### Expression of VMAT2 and DA metabolic pathways in human astrocytes

In order to assess the potential translational relevance of our findings, we obtained frontal brain cortex samples from human subjects and used previously validated polyclonal antibodies raised against tyrosine hydroxylase (TH), catechol-o-methyl transferase (COMT), MAOB and VMAT2 (50) to check for the expression of *bona fide* proteins involved in the synthesis, storage and degradation of dopamine in astrocytes. In line with previous findings in human and rodents brain tissues (50,63,64), immunolabelling experiments did not detect any signal for TH, but it did reveal the presence of COMT, MAOB and VMAT2 in >80% of GFAP-positive cells (Supplementary Figure S5a). In particular, the VMAT2 signal was readily recognisable mainly in one of the previously identified four major morphologic sub-classes of GFAP-immunoreactive cells of adult human frontal lobe (65): large, typically tortuous and highly branched protoplasmic astrocytes with primary processes located in layers 2-6 (Figure 5a-c). Interestingly, the signal was virtually absent or weak in both human and mouse GFAP-positive cells of the visual cortex (Supplementary Figure S5b), thus further validating the specificity of the staining and confirming the frontal cortex-specific expression of VMAT2 in astrocytes. We then checked for the presence of DA in the VMAT2-expressing GFAP-positive cells using a previously validated polyclonal antibody raised against dopamine (50) (Supplementary Figure S5c). In line with findings in rodents (Figure 5c-d, mouse), immunostaining showed the co-localisation of VMAT2 and dopamine in human astrocytes located in all of the layers of human cortical brain tissue (Figure 5e), thus indicating that astrocytes expressing VMAT2 and dopamine are also present in human cortical tissue.

**Figure 5.**
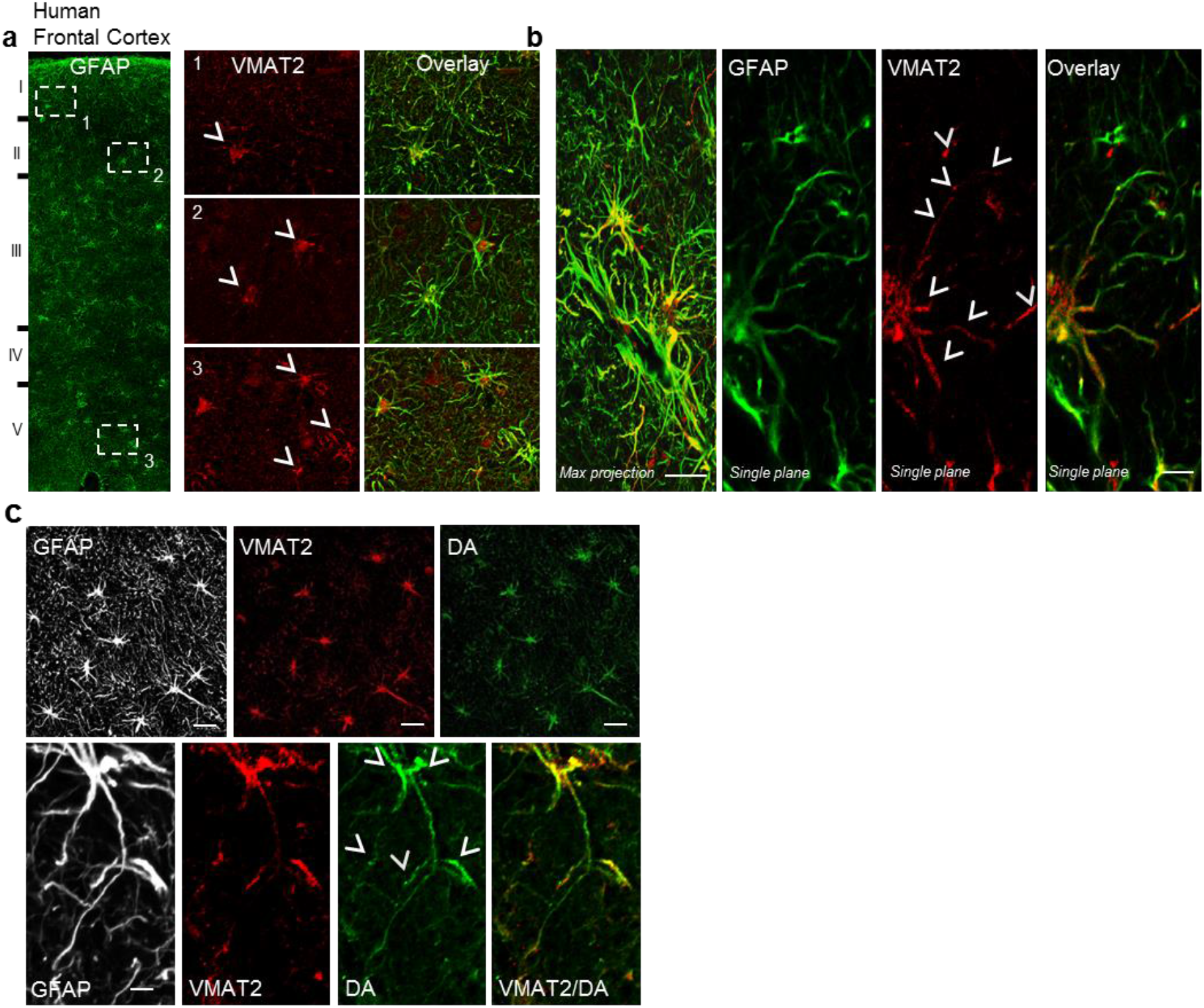
Expression of VMAT2 in astrocytes of the human brain frontal cortex. All images (representative of six slices from two tissue samples) are z projections of stacks 8-μm thick. (**a-b**) The expression of VMAT2 (red) in the cell bodies and processes of astrocytes identified by the cytoplasmic marker GFAP. Some astrocytes exhibit a high VMAT2 labeling of the soma (arrowheads, high magnification in a), whereas in others, VMAT2 labeling is mainly expressed on processes (arrowheads, high magnification in b). Bar, 30 μm (**c**) The expression of VMAT2 and dopamine in the cell bodies and processes of astrocytes identified by the cytoplasmic marker GFAP. Bar, 30 μm

## Discussion

We provide evidence that VMAT2 is expressed in human astrocytes located in the frontal cortical regions and that adult mice lacking VMAT2 in PFC astrocytes show abnormal grooming behaviours and increased anxiety. This phenotype is potentially interesting because it may be related to the aetiology of human OCD spectrum disorders such as human trichotillomania, which is characterised by repetitive stereotypical hair-pulling from various sites and leads to noticeable hair loss (66).

Rodent self-grooming is now widely recognised as a suitable behavioural output for studying the neural circuits underlying the generation of complex patterns of repetitive behaviours that may be relevant to human psychiatric disorders (4) and, over the last ten years, it has been suggested that some transgenic mice characterised by aberrant repetitive grooming behaviour can be used as models of OCD or OCD spectrum disorders. The grooming behaviour of VMAT2-deficient mice has some similarities to that identified in mice lacking synapse-associated protein 90/postsynaptic density protein 5-associated protein 3 (SAPAP3-/-) (67), which essentially consists of a pathologically increased number of short grooming sessions. The complex sequenced structure of mouse self-grooming is characterised by repeated stereotyped movements or “syntactic chains” (68) that start of a series of elliptical bilateral paw strokes near the nose (paw and nose grooming) and a series of unilateral paw strokes from the mystacial vibrissae to below the eye (face grooming), followed by a series of simultaneous bilateral strokes backwards and upwards (head grooming) and body licking. The duration of a single VMAT2-deficient mouse grooming event varies widely, but more detailed analysis revealed a deficit in grooming syntax, with many shorter (<3-second; data not shown) grooming events appearing by early adulthood (about P40) that seem to consist of two phases: paw and nose grooming and face grooming. The young onset of this aberrant repetitive behaviour suggests that the detection of hair loss at the front of the snout in later adulthood is due to this deficit in grooming syntax. Our observations are compatible with the early life appearance of aberrant grooming behaviour in mouse models of OCD (58) and patients with OCD and trichotillomania (58,69).

Analyses of rodent grooming behaviour have suggested that pattern-generating signals organise the physical movements made during individual grooming bouts (3,70), and multiple brain regions such as the basal ganglia, striatum (ST) and various cortical areas seem to be involved in implementing the syntactic chain (3,70). The execution of motor behaviour is also controlled by top-down projections from cortical areas (including the PFC) to the dorsal ST that form strong excitatory connections with striatal neurons (71,72). It is believed that these cortico-striatal connections make up a neural system that is required for habit learning, the routine performance of habits, and the acquisition of stereotype behaviour, and the abnormal functioning of this circuitry has been linked to the abnormal repetitive behaviour of patients with OCD-spectrum disorders (73).

We have shown VMAT2 expression in astrocytes of the frontal and prefrontal cortex, brain areas that are associated with rodent grooming behaviour and OCD circuitry. Immunohistochemical analyses of human and rodent cortical tissue show that VMAT2 is enriched in astrocytes of the frontal cortex but undetectable in astrocytes of the visual cortex, thus suggesting that astrocyte VMAT2 plays a functional role in frontal cortical regions. VMAT2-labelled astrocytes are a sub-population of cortical astrocytes (i.e. those located in frontal cortical regions), but the fact that we found that selective inactivation of VMAT2 in the mPFC is sufficient to induce pathological OCD-related behaviour favours the hypothesis that the different astrocyte populations in the brain play distinctive roles. However, the cell mechanisms regulating VMAT2 expression in specific astrocyte sub-populations require further investigation.

The mechanism underlying the way in which astrocyte VMAT2 affects behaviour may depend on the role of the vesicular transporter in regulating extra-cellular DA levels (50). In line with this hypothesis, we found that the re-expression of VMAT2 in the PFC sub-population of astrocytes and systemic treatment with L-DOPA, which has been reported to restore DA levels in the PFC of aVMATcKO mice (50), are sufficient to rescue abnormal grooming behaviour and anxiety. This provides a causal link between the absence of VMAT2 in astrocytes, decreased DA levels and the onset of OCD-like behaviour, and supports the hypothesis that increasing DA levels in the PFC may be therapeutically beneficial to patients with OCD.

The positive effect of L-DOPA on the abnormal grooming behaviour of VMAT2-deficient mice is due to the fact that L-DOPA restores extra-cellular DA levels in the PFC, and thus restores the normal basal activity of the excitatory circuits of aVMATcKO mice (50). The presence or absence of VMAT2 in mPFC astrocytes also seems to have a detectable electrophysiological effects. Previous measurements of basal synaptic transmission made by recording input/output (I/O) curves have shown that synaptic efficacy is significantly increased in the absence of astrocyte VMAT2, and that this is due to the lack of the tonic suppression of excitatory transmission caused by DA acting on D2 receptors (50). Our use of optogenetic projection targeting allowed us to assess mPFC cortico-striatal connections by measuring the ratio between post-synaptic AMPAR and NMDAR excitatory current amplitudes, and revealed an increase in the efficacy of synaptic transmission. Like its effect on excessive grooming behaviour, the re-insertion of VMAT2 in astrocytes of the mPFC was sufficient to rescue this potentiation of synaptic transmission, thus suggesting that the abnormal repetitive behaviour is associated with stronger mPFC-to-striatum transmission. Similar synaptic potentiation has been found in the orbitofrontal cortex-striatal connections of a mice model of compulsion in which the compulsive optogenetic self-stimulation of dopaminergic neurons is associated with peak activity in the terminals of projections from the orbitofrontal cortex (58), thus suggesting that compulsive behaviour in general may be explained by the failure of the top-down inhibition of stimulus–response associations attributed to the PFC (74,75).

Taken together, our findings not only indicate a novel role of PFC astrocytes in regulating cortico-striatal synaptic strength through DA homeostasis, but also show a phenotype that has not been previously associated with a decrease in astrocyte VMAT2. VMAT2 is an integral membrane protein that is expressed by aminergic cells in order to transport monoamines (particularly neurotransmitters such as DA, norepinephrine, serotonin and histamine) out of the cytoplasm and into the lumen of intra-cellular vesicles, including synaptic vesicles (76). By changing cytosolic and intra-vesicular monoamine concentrations, the up- or down-regulation of VMAT2 is crucial in regulating their homeostasis. Full VMAT2 knockout is lethal, and mouse studies have established that VMAT2 plays a critical role in maintaining catecholamine and serotonin levels in the central nervous system, and ensuring the availability of monoamines for exocytotic release from neurons (77,78). Given its crucial importance in regulating amines, human VMAT2 variants are very rare, but some have been associated with schizophrenia (79) and others with protection against alcohol neurotoxicity (80) Pathogenic variants in the gene encoding VMAT2 (i.e. *SLC18A2*) have only recently been described (81–83), cause severe forms of brain DA-serotonin vesicular transport disease and a variety of symptoms such as hypotonia, parkinsonism, tremor, developmental disability, and depression. Whether *SLC18A2* participates in the pathophysiology of OCD has never been investigated, but our findings indicate the existence of an OCD-like phenotype specifically caused by a knockout in this gene in astrocytes. The presence of VMAT2 in astrocytes has only recently been identified by means of transcriptome analysis and immunohistochemistry (50), and a novel mRNA splice variant of *Drosophila* VMAT (DVMAT-B) has been found in a small sub-set of the glial cells in the lamina of the fly’s optic lobe, where it regulates the homeostasis of histamine (84). Consequently, the pathological conditions associated with the modulation of this gene in astrocytes are still unknown and require further investigation.

## METHODS

### Maintenance, breeding and genotyping

All of the animal studies were approved by the *Service de la consommation et des affaires vétérinaries du Canton Vaud*, the Institutional Animal Care and Use Committee of the University of Geneva and the animal welfare committee of the Canton of Geneva, in accordance with Swiss law. The mice were group-housed with littermates in standard housing with a 12:12-hour light:dark cycle. The hGFAPcre^ERT2^ mice (85) and ROSA26-eYFP-hGFAPcre^ERT2^ mice (86) were obtained from Frank Kirchhoff (Molecular Physiology, University of Saarland, Germany), the VMAT2^Loop/Loop^ mice (53) from Bruno Giros (Douglas Mental Health University Institute, Canada), and the tdtomato^lox/lox^ mice (AI14, Jackson Lab) and Thy1-EGFP mice from Joshua R. Sanes (Harvard University, USA). The mice used had a C57BL/6 background.

The hGFAPcre^ERT2^ sequence was identified from phalange biopsies using the following primers: 5’- CAGGTTGGAGAGGAGACGCATCA-3’, 5’- CGTTGCATCGACCGGTAATGCAGGC-3’. The VMAT2^lox/lox^ sequence was identified using the following primers: 5’- GACTAGGGACAGCACAAATCTCC-3’, 5’-GAAACATGAAGGACAACTGGGACCC-3’. The ROSA26-EYFP sequence was identified using the following primers: 5’-AAAGTCGCTCTGAGTTGTTAT-3’, 5’-GCGAAGAGTTTGTCCTCAACC-3’, 5’-GGAGCGGGAGAAATGGATATG-3.

The PCR reaction product coupled with Syber green migrated in a 1.5% agarose gel, and the bands were revealed by UV light.

### Tamoxifen treatment

Tamoxifen (TAM, Sigma-Aldrich) was injected on P20; the treatment protocol has been previously described (50–52)

### Virus preparation

Self-inactivated (SIN) lentiviruses contain the central polypurine tract (cPPT) sequence, the mouse phosphoglycerate kinase I promoter (PGK), the woodchuck post-regulatory element (WPRE) sequence, and the target sequence of miR124 (62). A Gateway system (Invitrogen) was used to clone mouse VMAT2 and GFP cDNA, into the pENTR-D-TOPO plasmid (Invitrogen) and perform an LR clone reaction to transfer it into the destination lentiviral vector SIN-cPPT-PGK-Gateway-WPRE-miR124T plasmid. The viruses (lentiGFP, lentiVMAT2 and lentiCre) were produced in 293T cells using a four-plasmid system as previously described (87), and were pseudotyped using the G protein of mokola lyssaviruses (62).

### Stereotaxic intracranial injections

P25 mice were anesthetised using isoflurane at 5% (w/v), placed in a small animal stereotaxic frame (David Kopf Instruments), and maintained at 2.5% isoflurane (w/c) for the duration of surgery. Corneal and pinch reflexes were regularly tested in order to confirm the depth of the anesthesia. Lacryvisc (Aicon, Switzerland) was used to prevent corneal drying, and lidocaine was topically applied to the skin overlying the skull. After exposing the skull under aseptic conditions, a small hole was drilled into the skull overlying the prefrontal cortex (AP + 2.0 mm, L ± 0.2 mm and DV −2.0 mm) [Sohal et al., 2009]. LentiVMAT2, lentiCre and/or lentiGFP were injected (1μl total volume) bilaterally through a Hamilton syringe at a rate of 100 nl min-1 using a CMA400 Pump (CMA System). Adeno-associated virus AAV8-hSyn-Chrimson-GFP (0.8 μL total volume; Duke University, Durham, USA) was injected at P30 into control LoxTAM and aVMAT2cKO mice bilaterally using a thin glass pipette connected to an injection wheel. After the surgical procedures, the mice were returned to their home cages at least for two weeks in order to allow maximum gene expression.

### Tissue preparation, immunohistochemistry and histology

The mice were deeply anesthetised using intraperitoneal sodium pentobarbitone (6 mg/100g body weight) and immediately perfused intracardiacally with fresh 4% paraformaldehyde in 0.1 M phosphate-buffered saline (PBS, pH 7.4). Their brains were post-fixed overnight, equilibrated in 30% sucrose at – 20°C using a cryostat (Leica), and stored at −80°C.

Brain sections were permeabilised for 45 minutes in phosphate-buffered saline containing 0.3% Triton X-100, and 15% donkey or goat serum, and then immunolabelled overnight at 4°C using the following primary antibodies: mouse-GS (Chemicon, 1:1000) (88), rabbit-GFP (Chemicon, 1:200) (89), rabbit-GFAP (Chemicon, 1:1000), rabbit-OCT3 (Alpha Diagnostics, 1:100) (63), mouse-dopamine (Millipore, 1/100), rabbit-dopamine (Millipore, 1/1500), mouse-COMT (BD-Transduction Lab, 1/1000) goat-MAOB (Santa Cruz, 1:100), rabbit-VMAT2 (Synaptic System, 1:5600; Chemicon, 1:1000), and rabbit-TH (Millipore, 1:1000) (90). The day after incubation with primary antibodies, the brain sections were washed three times in PBS for 10 minutes and then incubated for 1.5 hours at room temperature with fluorescent secondary antibodies (AlexaFluor, Invitrogen Molecular Probes, Eugene, Oregon, USA: goat anti-mouse 488, 555, and 633; goat anti-rabbit 488, 555, and 633; and donkey anti goat 488; 1:300 and 1:400) diluted in PBS. Finally, the nuclei were counterstained with 4’, 6-diamidino-2-phenylindole (DAPI) (Invitrogen, 1:10000) and then washed before mounting with FluorSave (Calbiochem).

The brain was extracted and washed in PBS. Fragments of normal structure from human brain temporal cortices were removed as part of the planned margin of resection surrounding a neoplastic lesion from patients operated at the Section of Neurosurgery in the Policlinic San Matteo. Surgery was performed according to the recommendations of the Institutional Review Boards and in full agreement with the Declaration of Helsinki. Samples were fixed in 4% formaldehyde for 24 h and then washed in PBS. Human subjects samples from informed and consenting human patiens were collected before 2011 as previously described (91).

All images were collected on a Leica confocal imaging system (TCS SP5) with a 40x (1.4 NA) or 63x (1.4 NA) oil immersion objective. Sections were acquired every 0.4 μm, and the confocal images were analysed using Imaris 7.6.3 (Bitplane AG, Zurich, Switzerland) or Adobe Photoshop CS5 software (Adobe System Inc., San José, California, USA).

### *In vivo* microdialysis

The mice were anaesthetised with isoflurane and placed in a stereotaxic frame using a mouse adaptor (David Kopf Instruments) with modified ear bars. Microdialysis probes were implanted in the PFC at the following coordinates relative to the bregma: AP +2.0 mm, ML +0.5 mm and DV −3.0 mm, with the tooth-bar set at 0 mm. The active dialysis surface length of the membrane was 2 mm. The probe was secured in place using dental cement on the skull.

The microdialysis experiments started 24 hours after surgery. Ringer solution (125 mM NaCl, 2.5 mM KCl, 1.26 mM CaCl2, 1.18 mM MgCl2, 0.20 mM NaH2PO4) was perfused through the microdialysis probe at a flow rate of 1.0 μl/min using a high precision pump (CMA 400 syringe pump, CMA Sweden) (50, 91). The experiments were performed during the light period of the light/dark cycle, and the mice were tested in their home cages. After an equilibration period of at least two hours, the dialysates were collected into small Eppendorf tubes containing 11.7 μL of acetic acid every 30 minutes, and stored at −69°C until high-performance liquid chromatography (HPLC) analysis.

Dopamine levels were quantified by means of HPLC with electrochemical detection as previously described (50) with some modifications. The samples were injected into an MD-150 column (3 M, 3.2×150 mm, Thermo Fisher Scientific,) using a Thermo Scientific Dionex Ultimate 3000, and dopamine was detected at 32°C using an ECD-3000RS electrochemical detector (Thermo Fisher Scientific) set at a potential of 250 mV against an Ag/AgCl reference electrode. The signal was analysed using Cromeleon, software. The mobile phase was 75 mM sodium dihydrogen phosphate monohydrate, 1.7 mM ottane sulphonic acid sodium, 100 M triethylamine (TEA), 25 M EDTA, 10% acetonitrile, pH 3.00 with phosphoric acid. The flow rate was 0.5 mL/min.

### Morphological analysis of dendrites and spines

Dendritic spine density and spine morphology was assessed as previously described (50,91). The spine analysis was made using two fluorescent transgenic mice strains: control LoxTAM-Thy1EGFP and VMAT2^loxp/loxp^-Thy1EGFP mice obtained by crossbreeding Thy1EGFP with VMAT2^loxp/loxp^ mice. Confocal microscopy of post-fixed slices was performed using a Leica confocal imaging system (TCS SP5) with a 40× (1.8 NA) or 63x (2.8 NA) oil immersion objective. The mPFC field between Bregma coordinates 2.34 mm and 1.7 mm was analysed, with 50 μm brain sections being acquired every 0.4 μm. The number of spines on 10-20 neurons per mouse was counted using image J software, and spine density was expressed as the number of spines divided by dendritic length (92).

### Patch-clamp recordings

The mice were anesthetised with isoflurane (5%) and decapitated, and their brains were quickly removed. Brain slices (220 μm thick) were cut using a vibratome in ice-cold oxygenated artificial cerebrospinal fluid (aCSF) containing NaCl 119 mM, D-glucose 11 mM, NaHCO3 26.2 mM, KCl 2.5 mM, MgCl_2_ 1.3 mM, NaH_2_PO_4_ 1 mM, and CaCl_2_ 2.5 mM. The slices were re-covered in aCSF at 30°C for 15 minutes, and then kept at room temperature until use. The recordings were made in aCSF with 100 μM picrotoxin at 30°C. The internal solution consisted of 130 mM CsCl, 4 mM NaCl, 5 mM creatine phosphate, 2 mM MgCl_2_, 2 mM Na_2_ATP, 0.6 mM Na_3_GTP, 1.1 mM EGTA, 5 mM HEPES, 5 mM QX-314, and 0.1 mM spermine. The currents were amplified using Multiclamp 700B (Axon Instruments), filtered at 2.2 Hz, and digitalized at 20 Hz. No correction was made for liquid junction potential (−3 mV). Cells with more than 20% change in access resistance were discarded. The light-evoked currents were induced at 0.1 Hz (1 ms pulse, 593 nm LED). The A/N ratio was calculated as the amplitude of the AMPA receptor currents recorded at −70 mV divided by the amplitude of the trace at +40 mV 20 ms after the peak. In some recordings, the NMDA receptor currents were blocked using 50 μM D-2-amino-5-phosphonopentanoate (AP-5). The NMDA receptor traces were computed by subtracting the traces recorded at +40 mV in the presence of AP-5 from the traces recorded at +40 mV in the absence of AP-5. The rectification index was calculated as chord conductance at −70 mV divided by chord conductance at +40 mV.

### Behavioural protocols

#### Open field test

The diameter of the round area used for the open field test was 70 cm, and the central zone line was 25 cm from the edge. The mice were placed in the centre of the area at the beginning of the test, and their movements were video-recorded and subsequently analysed using ANY-maze 4.7 (Stoelting).

#### Elevated plus-maze

The elevated plus-maze consisted of two open arms, two closed arms, and a central area elevated to a height of 50 cm above the floor. Mice were placed in the central area and allowed to explore the space for eight minutes.

#### Repetitive behaviours

The mice were observed for 10 minutes in their home cages equipped with fresh bedding, and the time spent in repetitive behaviours (grooming, digging and jumping) was measured. Grooming was defined as stroking or scratching of the face, head or body with the two forelimbs, or licking the body; digging was defined as the co-ordinated use of both forelimbs or hind legs to dig out or displace bedding materials; jumping was defined as rearing on the hind legs at the corner or along the side walls of the cage, and jumping in such a way as both hind legs are simultaneously off the ground.

### Statistical analysis

**A**ll of the analyses were made using GraphPad Prism 8.0.2 software and one-way analysis of variance (ANOVA) followed by Bonferroni’s or Tukey’s *post hoc* test. An unpaired t-test was used for two-sample comparisons. Two sample comparison were made with unpaired t-test if data was normally distributed, and with a parametric test if data was not normally distributed. Three groups were analysed with one-way ANOVA followed by Tukey post-hoc test. All of data are expressed as mean values ± the standard error of the mean (SEM).

