## Supplementary Figures for "Astrocytic VMAT2 in the developing prefrontal cortex is required for normal grooming behavior in mice"

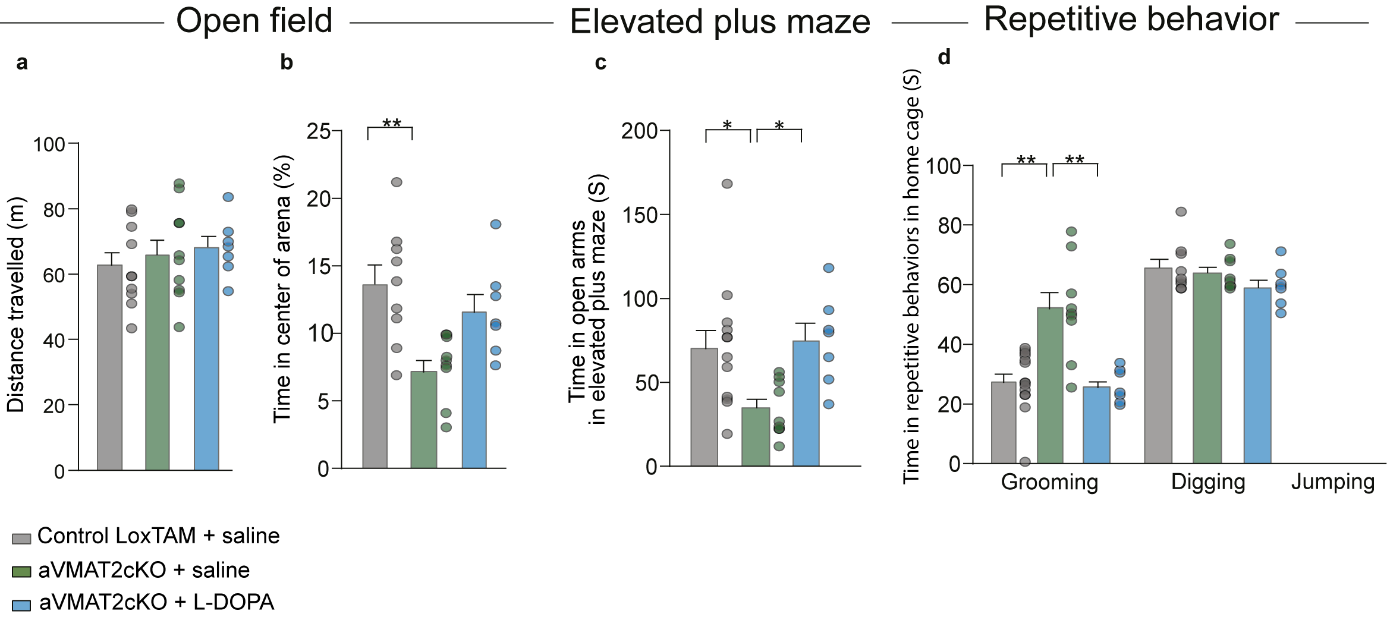

**Supplementary Fig 3. Chronic treatment with L-DOPA rescues the behavioural deficits of aVMAT2cKO mice. (a-b)** Open field tasks. (**a**) The histograms show the average total distance travelled by control LoxTAM and aVMAT2cKO mice treated with saline (grey and green) or L-DOPA (light blue) during 10 minutes of free exploration of a maze, and (**b**) the average percentage of time they spent in the center of the arena. **p<0.005 (n=8-9 in each group, one-way ANOVA, followed by Tukey’s post hoc HSD test). (**c**) Elevated plus maze tasks. The histograms show the average percentage of time that control LoxTAM and aVMAT2cKO mice treated with saline (grey and green) or L-DOPA (light blue) spent in the open arms. *p<0.01, (n=10-12 ; one-way ANOVA, followed by Tukey’s post hoc HSD test). (**d**) Repetitive behavioural tasks. Average time that control LoxTAM mic (grey), aVMAT2cKO mice treated with saline (green) and aVMAT2cKO mice treated with L-DOPA (light blue) spent in grooming and digging. **p<0.005 (n=8 each group; one-way ANOVA, followed by Tukey’s post hoc HSD test).

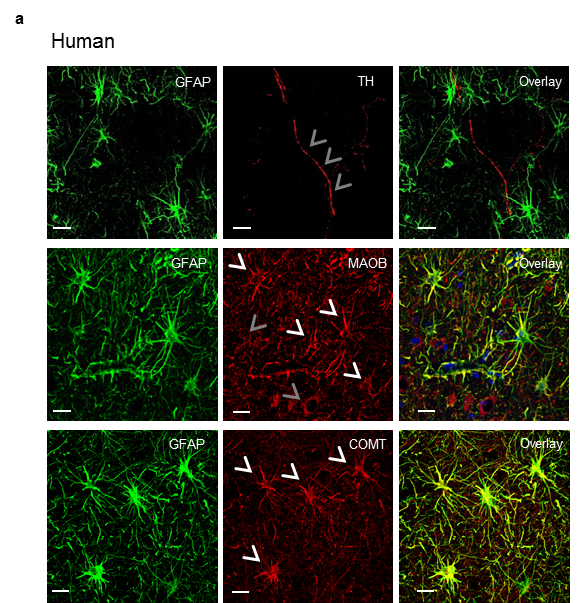

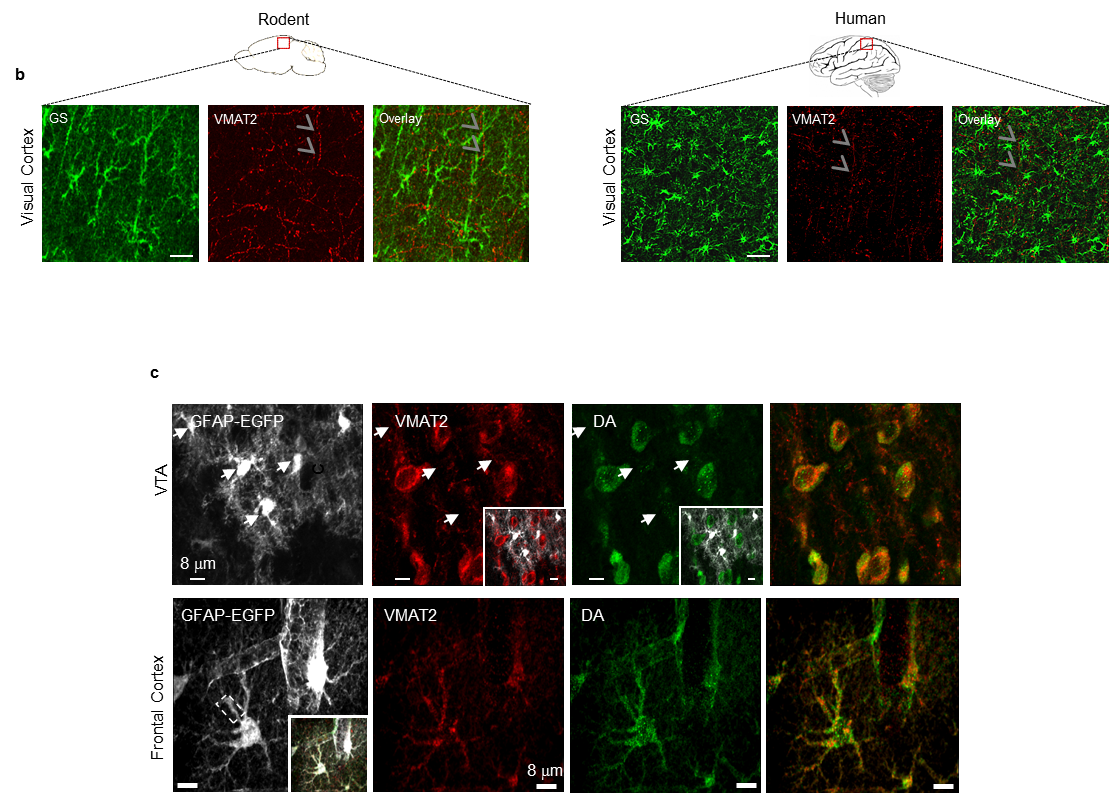

Supplementary Fig 5. Expression of VMAT2 expression of *bona fide* proteins involved in the synthesis, storage and degradation of dopamine in astrocytes of the human brain frontal cortex. All images (representative of six slices from two tissue samples) are z projections of stacks 8-µm thick. (**a**) The expression of TH, MAOB and COMT (red) in the cell bodies and processes of astrocytes identified by the cytoplasmic marker GFAP. Note the absence of TH immunostaining in astrocytes. Bar, 30 µm (**c**) The absence of expression of VMAT2 in astrocytes located in the visual cortex of rodent and human brain tissues. Astrocytes have been identified by the cytoplasmic marker glutamine synthase (GS) and Bar, 30 µm. (**d**) The expression of dopamine and VMAT2 in dopaminergic neurons of the ventral tegmental area (VTA) and in astrocytes of the prefrontal cortex (PFC) in rodent tissues. Bar, 8 µm.
